# Structural Basis for RNA-guided DNA degradation by Cas5-HNH/Cascade complex

**DOI:** 10.1101/2024.07.22.604555

**Authors:** Yanan Liu, Lin Wang, Qian Zhang, Pengyu Fu, Lingling Zhang, Heng Zhang, Hongtao Zhu

## Abstract

Type I-E CRISPR (clustered regularly interspaced short palindromic repeats)–Cas (CRISPR-associated) system is the most widely investigated RNA-guided adaptive immune system in prokaryotes against the foreign genetic elements. Unlike the previously characterized Cas3 nuclease in the typical Type I-E system that exhibits uncontrolled progressive DNA cleavage, a recently discovered HNH domain linked to the Cas5 subunit functions as the nuclease for precise DNA cleavage, highlighting the potential of this new Type I-E variant system as a precise genome editing tool. Here, we present five near-atomic cryo-EM (cryo-electron microscopy) structures of *Candidatus Cloacimonetes bacterium* Cas5-HNH/Cascade complex either bound or unbound to DNA. We revealed that the HNH domain extensively interacts with the adjacent subunits, including Cas6, Cas8 and Cas11. The related mutations of the crucial identified interactions can significantly weaken the performance of the enzyme. Upon the binding of DNA, the Cas-HNH/Cascade complex adopts a more compacted conformation with the subunits moving towards the center of the nuclease, thus activating the nuclease. We further identified four conserved cysteines that form a zinc-finger structure in the HNH domain and the mutations of these four cysteines can totally abolish the enzyme activity. Interestingly, we also discovered that the divalent ions such as zinc, cobalt and nickel down-regulate the enzyme performance by decreasing the stability of the Cascade complex. Together, our findings provide first structural insights into the assembly and activation of the Cas5-HNH/Cascade complex, opening new avenues for engineering this system for precise genome editing.

## Introduction

Prokaryotes utilize an adaptive immune system called CRISPR (clustered regularly interspaced short palindromic repeats)–Cas (CRISPR-associated) system to protect them from foreign genetic elements such as invading bacteriophage and plasmids^1–3^. CRISPR array is composed of a series of conserved repeats that are separated by variable spacer sequences which are adopted from alien nucleic acids^1,4–6^. Transcripts from CRISPR loci are processed into mature and short CRISPR RNAs (crRNAs), which assembles with Cas gene-encoded proteins into large ribonucleoprotein surveillance complexes^7,8^. The coevolution of bacteria with the invading viruses and other invasive gene components has resulted in remarkably diversified CRISPR-Cas systems^9,10^. CRISPR-Cas systems are now classified into two major classes: Class 1 and Class 2, which can further be divided into 6 types (type I to VI) and numerous subtypes (e.g. type I-A to I-F)^11,12^. The Class 1 system that used by most bacteria and archaea for their surveillance, is featured with multiple effectors^13^. However, in Class 2 systems, a single protein such as Cas9, Cas12 or Cas13 serves as the effector, and this property allows Class 2 systems to be widely used in biotechnological fields as genome editing tools^14–17^.

Canonical Cascade (CRISPR-associated complex for antiviral defense) surveillance complex in type I-E systems contains five stoichiometrically unequal Cas proteins, including Cas5, Cas6, Cas7, Cas8 and Cas11^18–22^. In type I-E system, the mechanisms of bacteria immune reaction include three stages: spacer acquisition (also known as adaptation), processing of pre-crRNAs and interference^23,24^. During the first stage, Cas1 and Cas2, which form a butterfly-shaped dimer, are required to capture the short foreign genetic elements and integrate them into the host CRISPR array as the spacers, thus enabling the bacteria to gain new immune memories^25–27^. In the second stage, the endoribonuclease Cas6 recognizes the pre-crRNA and cleaves it into small matured crRNA, which has a 5’ handle, a spacer and a 3’ stem-loop structure with a length of approximate 61-nt^28,29^. After the maturation of the crRNA, Cas6 remains closely bound to the 3’ stem-loop of the crRNA, allowing the other Cascade members to be recruited to form a complete Cascade complex^30,31^. During the third stage, the Cascade complex identifies the invasive DNA by a protospacer-adjacent motif (PAM) and a complementary sequence to the crRNA^32,33^. And then, a nuclease and helicase protein known as Cas3 is recruited to digest the foreign DNA^34,35^. Though type I systems are characterized by Cas3, yet Cas3 is not a stable member in the Cascade complex^36,37^. Recently, a variant of type I-E system lacking Cas3 from the *Candidatus Cloacimonetes bacterium* was discovered using sophisticated bioinformatics based on fast locality-sensitive hashing-based clustering (FLSHclust) algorithm^38^. Interestingly, an HNH nuclease domain that replaces the role of Cas3 was fused to the C-terminus of Cas5^38^. However, the architectures and molecular basis of this Cas5-HNH/Cascade complex are unknown.

Here, we reported five cryo-electron microscopy (cryo-EM) structures of the immunological nucleoprotein complex Cas5-HNH/Cascade complex at near-atomic resolution, including three structures in the absence of the target DNA and two structures bound with the target single-strand DNA (ssDNA). These cryo-EM structures revealed structural insights into regulation and activation of the HNH nuclease domain, distinct from the previously known typical type I system. We also found that, after binding to the target ssDNA, the Cas5-HNH/Cascade adopts a compacted conformation compared with the complex without ssDNA. Moreover, we discovered a zinc-finger structure in HNH domain which plays important roles in regulating the function of Cas5- HNH/Cascade complex.

## Results and discussions

### Biochemistry and structure determination of Cas5-HNH/Cascade complex

The wild-type Cas5-HNH/Cascade complexes were recombinantly expressed in *E.coli* with a 10XHis tag on Cas8. The SDS-PAGE experiment was further used to confirm that all of the required subunits including Cas5-HNH, Cas6, Cas7 and Cas11 were co-purified by Cas8 (Supplementary Fig.1a). Prior to the cryo-EM studies, we first performed digestion assay experiments to validate the enzyme activity, the results justified that the wild type of Cas5-HNH/Cascade complex can digest the targeted DNA efficiently. Previous study has proved that Cas5-HNH/Cascade complex has the ability to digest both ssDNA and double strand DNA (dsDNA)^38^. In order to obtain the complex of Cas5-HNH/Cascade with target DNA, we utilized fluorescence-detection size-exclusion chromatography (FSEC^39^) to test if the target DNA binds the complex. We first performed the experiments in the presence of ssDNA. A fluorophore group FAM was added to the 3’ end of the ssDNA. Our FSEC data featured an obvious peak migration of ssDNA, indicating that ssDNA binds to Cas5-HNH/Cascade complex (Fig. 1j and Supplementary Fig.1b,c). Before loading the sample to the size exclusion column (SEC), the Cas5-HNH/Cascade complex was incubated with 61-nt of the single-stranded guide RNA and 60-nt target ssDNA. The SEC trace showed a mono dispersed sample that was obtained (Supplementary Fig.1a). We then subjected the complex to cryo-EM imaging and data analysis. Both the raw micrographs and the 2D class averages indicated a good quality of the cryo-EM dataset (Supplementary Fig.3). After the 3D analysis, two conformations were resolved including an intact complex (ssDNA_Conf1) and one partially assembled complex (ssDNA_Conf2), which were at an overall resolution of 3.00 Å and 2.81 Å, respectively (Fig.1b-g, Supplementary Fig.3 and Supplementary Fig.4a-f). The intact complex is well resolved with clear features of the side chains, allowing us to build reliable models (Supplementary Fig.5). However, we only resolved the density of target ssDNA which are paired with the crRNA (Fig.1h-i). We speculated that the missing density of the rest ssDNA is not defined probably due to flexibility.

**Fig 1.**
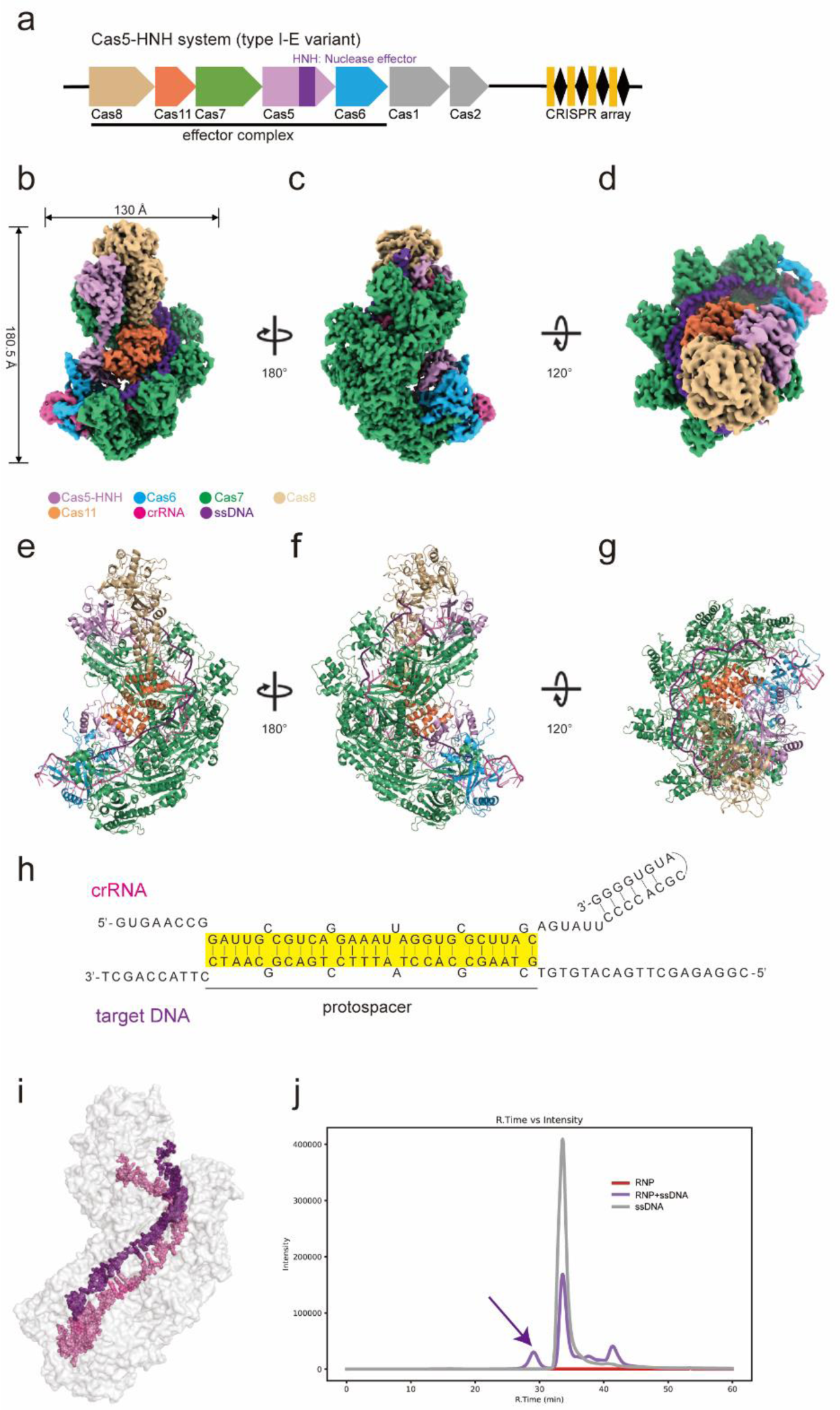
The Cryo-EM structures of Cas5-HNH/Cascade in Type I-E CRISPR-Cas system. **a,** Locus diagrams of Cas5-HNH system from *Candidatus Cloacimonetes* bacterium. **b-d,** Cryo-EM map of Cas5-HNH/Cascade shown in side views (**b,c**) and top down view (**d**). Six copies of Cas7 are colored in green. The main lobe of Cas5 and HNH domain are colored in plum. Cas6, Cas8 and Cas11 are colored in blue, burlywood and coral, respectively. The crRNA is shown in hot pink and the target ssDNA is colored in purple. **e-g,** Cartoon representation of the structure of Cas5-HNH/Cascade complex corresponding to (**b-d**). **h,** Schematic representation of crRNA and the target ssDNA. **i,** Target ssDNA (purple) and ssDNA (hot pink) was shown. The Cas5-HNH/Cascade complex are shown in surface representation. **j,** FSEC trace shows that wild type RNP binds with target ssDNA labeled by FAM. The arrow indicates the peak shifts of ssDNA.

Based on previous research, the canonical HNH motif is featured with three conserved important residues, including two histidine and one asparagine (Supplementary Fig.6c). In our structure, the first conserved histidine in HNH^Cas5^ (H310) is located at the end ofβ-strand 1 (Fig. 2a, Supplementary Fig.6a). Compared with the classic HNH motif, the second active site asparagine is replaced by an aspartate (D324) in HNH^Cas5^. Interestingly, the sequence alignment of type I-E Cas5-HNH family proves that the substitution of asparagine to aspartate is conserved (Supplementary Fig.6b-d), hinting the importance of aspartate in regulating the function of HNH^Cas5^. To further verify the importance of H310, D324 and H333, we constructed the mutants carrying H310A, D324A or H333A mutation. The biochemical analysis suggested that the alanine substitutions of H310 and D324 can totally impair the cleavage activity (Fig.2b). However, the third corresponding site H333A of HNH^Cas5^ has no influence in the nuclease activity (Fig.2b). Moreover, the electrophoretic mobility shift assay (EMSA) proved that the mutants have no influence of the binding of DNA (Supplementary Fig.10e). After obtaining the cryo-EM structures of Cas5-HNH/Cascade in complex with ssDNA, we attempted to capture the Cas5-HNH/Cascade complex without the ssDNA. After a thorough 3D classification for the cryo-EM data, three conformations were captured, including two intact complex and one partially assembled complex, which were at an overall resolution of 2.47 Å, 3.10 Å and 2.55 Å, respectively (Supplementary Fig.2, Supplementary Fig.4g-o and Supplementary Fig.7). And we thus named these three structures as apo_Conf1, apo_Conf2 and apo_Conf3.

**Fig 2.**
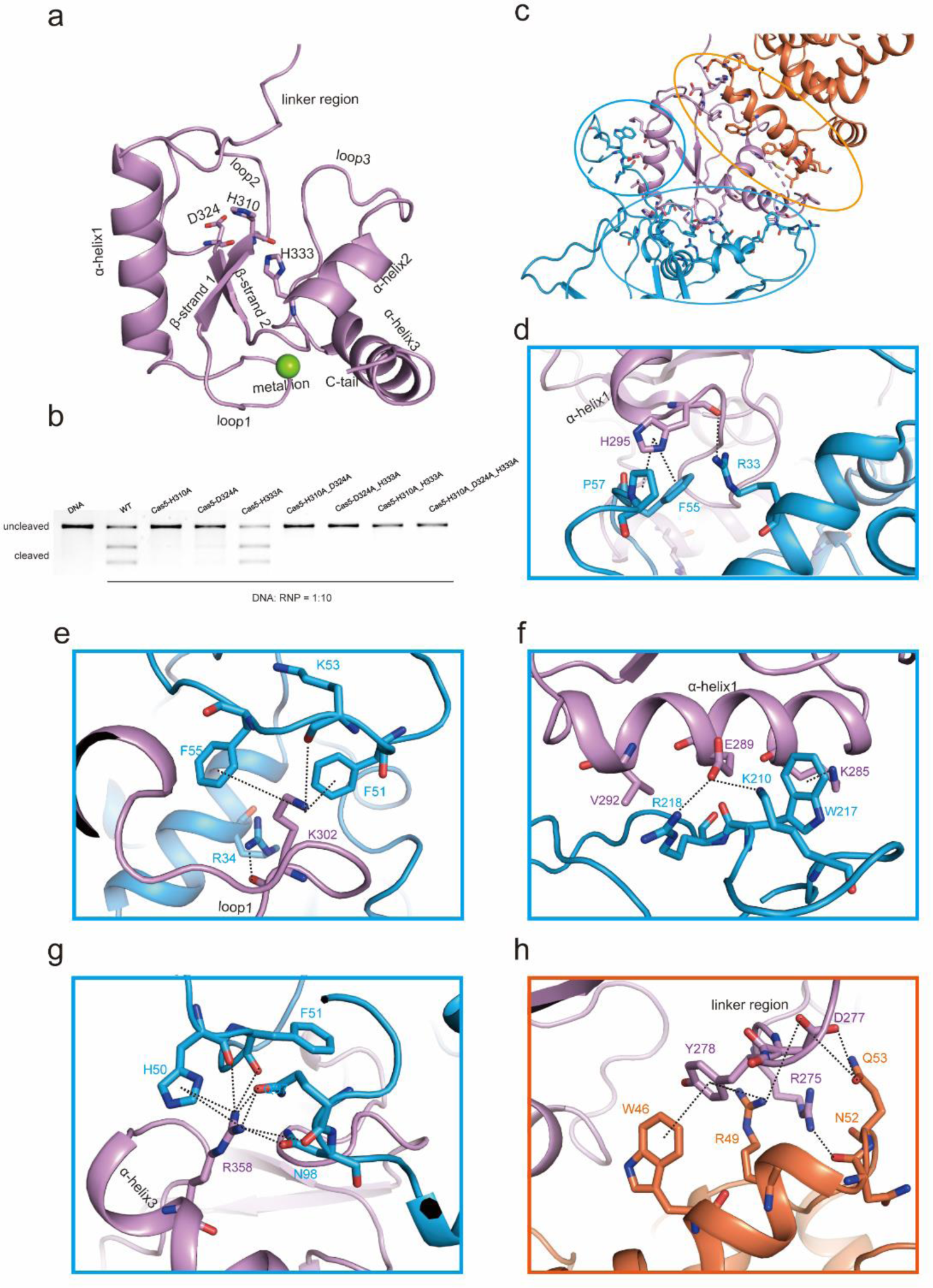
HNH domain functions as an endonuclease in type I-E variant Cas5-HNH /Cascade complex **a,** Structure of the HNH domain at the C-terminus of Cas5. HNH^Cas5^ is composed of three α-helices, two β-strands and several loops. Three featured residues (H310, D324, H333) are shown in sticks that were colored by elements. The divalent metal ion is shown by a green sphere. **b,** Plasmid digestion analysis results for the related mutants. **c-h,** Interactions between HNH domain and Cas6/Cas11 that are indicated by dashed lines. Residues of Cas5, Cas6, and Cas11 are colored by plum, blue and coral, respectively.

### Overall architecture of the Cas5-HNH/Cascade complex

The cryo-EM structures of Cas5-HNH/Cascade complex adopt a seahorse-shaped architecture. Take ssDNA_Conf1 for example, the width and length are 130 Å and 185 Å, respectively, which is roughly similar to these canonical Cascade complexes in type I-E CRISPR systems^18–22^ (Fig.1a-g). The length of apo_Conf2 is about 177.5 Å, 8 Å shorter than apo_Conf1 (184.7 Å, Supplementary Fig.7a-b). In the partially assembled complex ssDNA_Conf2/apo_Conf3, the HNH domain of Cas5, the N terminal domain of Cas6, Cas8 and Cas11 were not resolved. Due to the absence of Cas8, the height of apo_Conf3 is about 150 Å, much shorter than Conf1 and Conf2 (Supplementary Fig.7c). For the intact complex apo_Conf1 and apo_Conf2, six copies of Cas7 subunits, labeled as Cas7.1 to Cas7.6, packed along the crRNA to form a helical filament, which function as the scaffold of Cas5-HNH/Cascade complex (Supplementary Fig.8a). The measured distance of the center of masses (COM) between neighboring Cas7 subunits is approximate 30 Å (Supplementary Fig.11c-d). Though all of the Cas7 subunits adopt similar conformations, we still observed conformational differences among them (Supplementary Fig.8b). For example, we found that a β-hairpin in Cas7.1 subunit (residues 185-210) and several α-helices in Cas7.6 (residues 55-137 and 149-165) subunit are missing in our cryo-EM map (Supplementary Fig.8b). The ends of the Cas7 filament were capped by Cas8 and Cas6 respectively, while Cas5-HNH and Cas11 composed the back of the seahorse (Fig.1b-g). The 3’ crRNA handle forms a hook-like shape to clamp the Cas6 RNase, while 5’ end was covered by Cas5 (Fig.1b-g, Supplementary Fig.8c-f). Besides Cas7, we observed that RNA has wide interaction with Cas5, Cas6 and Cas8 through hydrogen bonds and π-π interactions (Supplementary Fig.9). For example, W160 and Y231 of Cas5 forms a π-π interaction with -7G and -5G, respectively. Meanwhile, R56 and K167 interacts with crRNA via hydrogen bonds (Supplementary Fig.9).

### HNH domain functions as nuclease in Cas5-HNH Cascade

The classic Cascade harbors two subtype-specific Cas11 subunit in its center^18–22^, but only one Cas11 subunit was observed in the Cas5- HNH/Cascade complex (Fig.1, Supplementary Fig.7, Supplementary Fig.8f and Supplementary Fig.11a-b). Instead, an HNH nuclease domain was fused to the C terminus of Cas5 (HNH^Cas5^) which was composed of two β-sheets and three α-helices (Fig.2a) which occupies the location of the second Cas11 subunit. Since Cas3 and HNH^Cas5^ both function as the nuclease, Cas3 is dispensable in the interference stage of bacterial immune response due to the presence of an HNH domain in type I-E system. After a structural alignment using DALI server, an HNH endonuclease in thermophilic bacteriophage GVE2 (GVE2 HNHE)^40^ got the highest score (PDB: 5H0M, Supplementary Fig.6b), hinting that the bacteria probably captured HNH domain from bacteriophage during the evolution.

The interactions related to HNH play critical roles in regulating the function of the complex. Investigation of the interfaces between HNH^Cas5^ and Cas6/Cas11 reveals wide interacts between them (Fig.2c). Hydrogen bonds which were involved H295, K302, and R358 in HNH^Cas5^, R33, R34, F51, K53, and N98 in Cas6 were observed (Fig.2d,e,g). At the same time, cation-π interactions were also found at the interface of Cas5 and Cas6, including K302^Cas5^-F51^Cas6^, K285^Cas5^-W217^Cas6^, and R358^Cas5^-H50^Cas6^ (Fig.2e-g). We also found salt bridges formed by E289^Cas5^ and R218^Cas6^/H220^Cas6^ (Fig.2f). To further validate the importance of these interactions, we designed multiple mutants and tested the efficiency of the enzyme. As expected, the mutations involved with the Cas5-HNH and Cas6 interfaces can significantly reduce the efficiency of DNA digestion (Fig.2b). Notably, mutation of E289A in HNH domain could disrupt the binding with target DNA (Supplementary Fig.10a,e). We also revealed that the mutant carrying M338A, V369A and F371A that locates in the last two helices of HNH domain, in addition to the mutants of their neighboring residues in Cas6, can impair its function (Supplementary Fig.10b,f).

Considering the HNH domain connecting the N-terminal domain of Cas5 via a long linker (Supplementary Fig.6a), we thus proceeded to validate if these amino acids located in the linker domain play roles in regulating the function of the nuclease. Interestingly, a combined mutation of R275A, D277A and Y278A in the linker domain resulted in a significant decrease in the nuclease activity (Fig.2h and Supplementary Fig.10d,h). Of note, these residues, including M338, F371, R275, D277 and Y278, are related to the interaction with Cas11 (Fig.2c,h). Taken together, our structural observations and biochemical experiments prove that the interactions between Cas6/Cas11 and HNH^Cas5^ are important for maintaining the function of the Cas5-HNH/Cascade complex. Mutations in these interfaces might bring conformational variations and consequently cause the disruption of the contacts which can further attenuate the cleavage of the DNA.

### Comparison of the structures reveals distinct conformational differences

A comparison between the complex bound with ssDNA and the unbound reveals obvious conformational differences (Fig.3). The global RMSD value between apo_Conf1 and ssDNA_Conf1 is 2.27 Å (Fig. 3d and Supplementary Fig.11c). We also observed that both ssDNA_Conf1 and apo_Conf2 adopt a more compacted conformation than apo_Conf1 (Fig.3a-c and Supplementary Fig.11c-f). The overlapped structures revealed that the major differences occur among subunits of Cas5, Cas8 and Cas7.6 (Fig.3d,e). Notably, after binding ssDNA, the HNH domain takes a shift towards Cas6 compared with apo_Conf1, yielding an RMSD value of 1.29 Å, especially at the region of α-helix 2 and 3 (Fig.3f).

**Fig 3.**
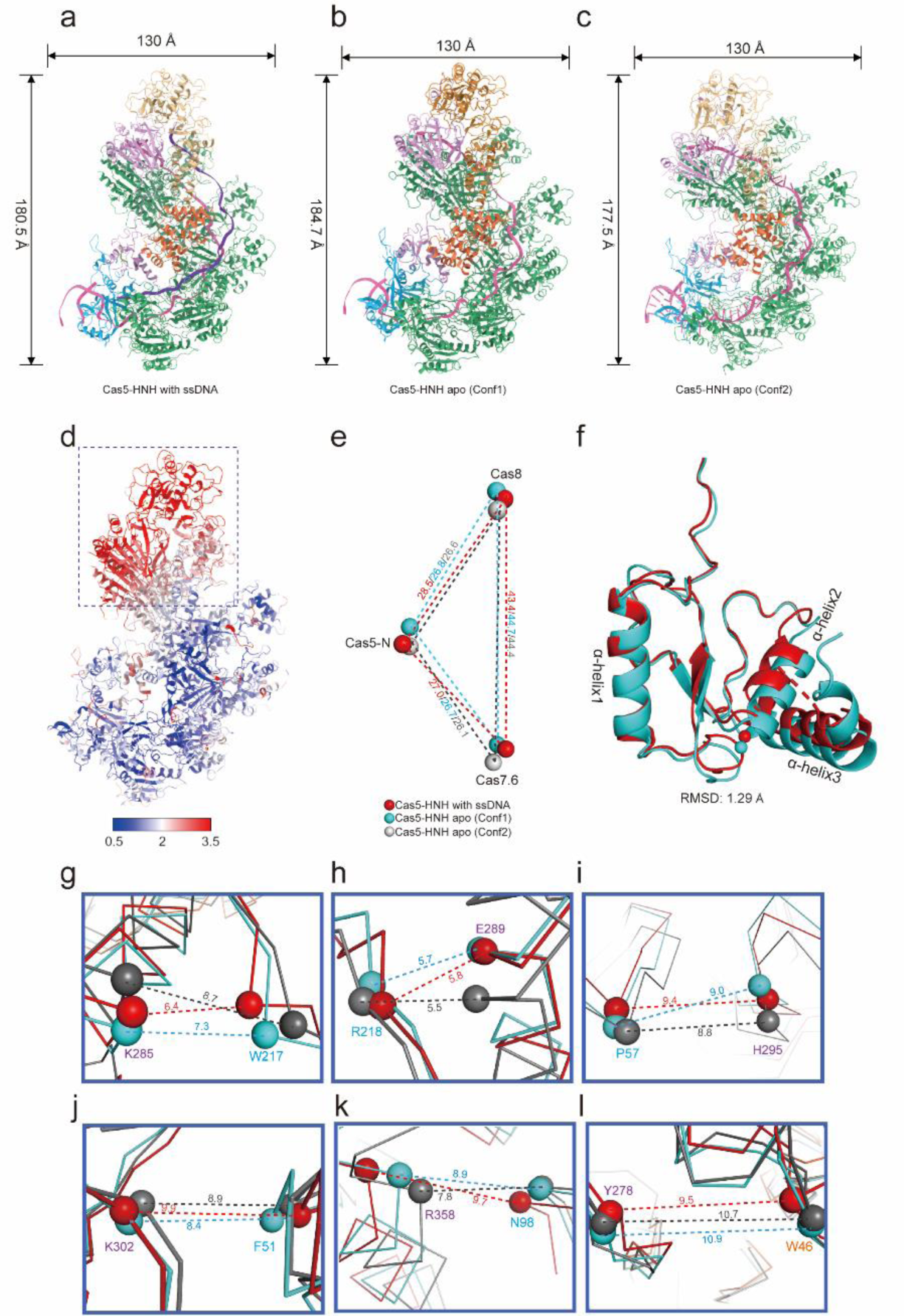
Comparison of the structures unbound or bound with ssDNA reveals distinct conformational differences **a-c,** Cartoon representation for Cas5-HNH/Cascade complex bound with ssDNA (**a**), apo_Conf1(**b**) and apo_Conf2 (**c**), respectively. The height and width are denoted in Å. **d,** RMSD distribution calculated between ssDNA_Conf1 and apo_Conf1. The major differences occur among subunits of Cas5, Cas8 and Cas7.6, which are indicated by dashed rectangle. **e,** Schematic illustrating the neighboring distances (Å) of centers of mass (COM) of Cas8, Cas7.6 and N-terminal domain (1-270) of Cas5. The Cas5-HNH bound with ssDNA, apo_conf1 and apo_conf2 are colored in red, cyan and grey, respectively. **f,** The overlap of the HNH domain from ssDNA_Conf1 (red) and apo_Conf1 (blue). The RMSD value is shown below the panel. **g-l,** The distances between the Cα atoms of representative amino acids on HNH^Cas5^. The color code is the same to panel (**e**).

**Fig 4.**
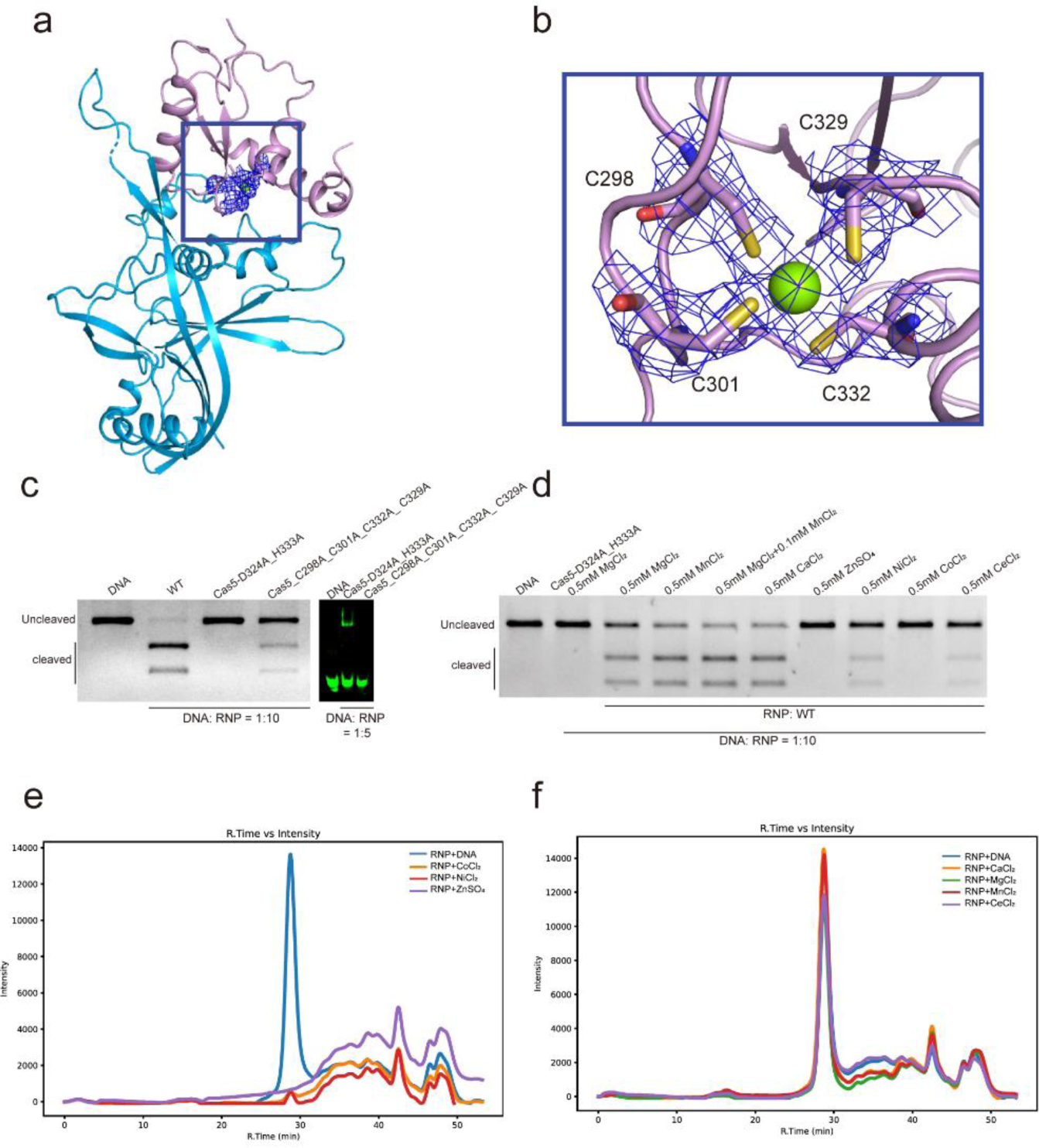
The zinc-finger like structure plays important roles in regulating the function of Cas5-HNH/Cascade complex **a-b,** The location of the four cysteines (C298, C301, C332, and C329) in HNH^Cas5^ was shown. Cas5 and Cas6 are colored in pink and blue, respectively. The boxed area in (**a**) is zoomed in panel (**b**). In panel (**b**), the cysteines are shown in sticks representation and the zoned cryo-EM density are shown in mesh representation. **c,** Biochemistry analysis shows that the mutation of these four cysteines will abolish both the binding and the digestion of target DNA. **d,** Biochemical analysis for the enzyme activities in the presence of metal ions including magnesium, manganese, calcium, cesium, cobalt, nickel and zinc. **e**, FSEC traces for the Cas5-HNH/Cascade complex in the presence of Co^2+^, Ni^2+^ and Zn^2+^. **f**, FSEC traces for the complex in the presence of 0.5 mM Mg^2+^, Mn^2+^ and Ca^2+^. The wavelength of the FSEC detector is 280nm.

The crRNA adopts significant conformations among the structures. We focused on the comparison associated with crRNA from these three intact conformations. The overlap of crRNA from apo_Conf1 and apo_Conf2 shows a clear shift nearby Cas7.5 with the calculated RMSD value of 2.28 Å (Supplementary Fig.11f). Interestingly, alignment of crRNA from ssDNA_Conf1 and apo_Conf1 represents a clear movement occurs at the two-end of the crRNA (Supplementary Fig.11g). In addition, the comparison indicates that the conformation of crRNA in apo_Conf1 and apo_Conf3 show high similarities to each other (Supplementary Fig.11h).

To explore the conformational changes triggered by the binding of target DNA, we aligned the crRNA in apo_Conf1, apo_Conf2 and ssDNA_Conf1 following the measurement of the shifts of the COM of the associated subunits (Supplementary Fig.11c-d). Previous research has proved that the activation of the CRISPR Cascade complex will undergo the movement of the subunits towards the nuclease center^20,22,35^. Consistently, we found that, upon the binding of ssDNA, the N terminal of Cas5, Cas7.6, and Cas8 moved towards the HNH^Cas5^. The calculated RMSD between ssDNA_Conf1 and apo_Conf1 for these three subunits is around 3.50 Å. We also measured the distances between the Cα atoms of these key amino acids on HNH^Cas5^ and their interaction partners (Fig.3g-l). Noted that, in ssDNA_Conf1, some of the distances between the key amino acids are closer to each other than the corresponding distances in apo_Conf1 (Fig.3g,l), while the other residues show further distances than apo_Conf1 (Fig. 3h,i,j,k). We also tested the mismatch tolerance of Cas5-HNH/Cascade complex using various mismatched DNA substrate. Mismatches at PAM-distal positions (29-32) did not affect the Cas5- HNH nuclease activity (Supplementary Fig.12). However, Cas5-HNH/Cascade complex could not tolerate the mismatch of PAM-proximal positions (19-17) (Supplementary Fig.12), which indicates Cas5-HNH/Cascade complex as a potential gene-editing tool.

### Divalent ions play important roles in regulating the cas5-HNH/Cascade complex

According to the sequence alignment (Supplementary Fig. 6c,d), we found that the HNH^Cas5^ is conserved among several species. By aligning the sequences, we discovered four conserved cysteines C298, C301, C329 and C332 in HNH^Cas5^. Many structures have demonstrated that such kind of four cysteines usually form a zinc-finger like structure. Interestingly, we observed strong connections among these four cysteines in our cryo-EM map (Fig.4a-b). As shown in Fig.4b, the density contributed by these four cysteines was featured with a cross shaped density, indicating a divalent ion probably coordinated here. To explore the roles these cysteines pairs playing in Cas5- HNH/Cascade, we constructed the mutants carrying four cysteines to alanine mutations. Then, we tested the efficiency of this mutant cleaving the target DNA. The results demonstrated that the mutation of these four cysteines would abolish both the binding and digestion of the target DNA (Fig.4c). The zinc finger like structures prompt us to investigate which metal ions that can regulate the function of Cas5-HNH. We tested several different kinds of metal ions, including magnesium, manganese, calcium, cesium, cobalt, nickel and zinc. First, we checked if these ions can modulate the enzyme efficiency of the cas5- HNH/Cascade complex. The results proved that, in the presence of 0.5 mM magnesium, manganese and calcium, the efficiency of Cas5-HNH/Cascade complex is comparable to each other (Fig.4d). However, when adding cesium, cobalt, nickel and zinc to the complex, the efficiency of the DNA digestion was significantly reduced. Especially for zinc and cobalt, the activity of the enzyme was nearly abolished (Fig.4d). To our surprise, when we performed the FSEC^39^ experiments in the presence of these metal ions, we found that the addition of cobalt, nickel and zinc can cause the disassociation of the complex (Fig.4e), and this may explain why these ions down regulated the efficiency of the enzyme. Though our results show that, in the presence of cesium can reduce the efficiency of the enzyme, yet it cannot trigger the disassociation of the complex (Fig.4f). Our evidence proved that these four cystines play important roles in maintaining the stability of the complex but also affect the activity of the complex, probably via binding the different kinds of divalent ions to modulate the function.

## Summary

In this study, we elucidated five near atomic structures of the newly discovered Cas5-HNH Cascade complex. Our structures show that the HNH^Cas5^, which cleaves the substrate DNA, replaced the Cas3 nuclease in the classic Type I-E CRISPR system. We revealed that the HNH^Cas5^ has extensive interactions with its nearby subunits including Cas6 and Cas11, and these amino acids associated with the interface of HNH^Cas5^ can bring side effects of the enzyme activity. The structural analysis revealed that, upon the binding of target DNA, the structure of Cas5-HNH/Cascade complex adopts a more compacted conformation, featured with the subunits Cas5, Cas7.6, Cas7.5 and Cas8 moving towards the nuclease center thus activating the enzyme. Interestingly, we found that four conserved cysteines in HNH^Cas5^ play important roles in regulating the function of the Cas5-HNH/Cascade. The mutation of cysteine to alanine can completely abolish the activity of Cas5-HNH/Cascade complex. We also uncovered that the Cas5-HNH/Cascade complex is vulnerable to divalent ions including cobalt, nickel and zinc which can cause the disassociation of the complex, and consequently down regulates the efficiency of the enzyme. In contrast, the enzyme can benefit from divalent ions like magnesium, manganese and calcium. Together, our structures can provide significant details for understanding the function of Type I-E CRISPR-Cas system.

## Methods

### Protein expression and purification

The Cas genes encoding full-length Cas5-HNH, Cas6, Cas7, and Cas8 proteins of the type I-E Cas5-HNH system is from *Candidatus Cloacimonetes bacterium ADurb.Bin088*. Cas5-HNH and N-terminal His_10_-tagged Cas8 were fused into a pCDFDuet vector. Cas7 and Cas6 genes were cloned into the pRSFDuet-1 vector, and the gene encoding a 61-nt crRNA was inserted into a pACYCT2 vector. All plasmids were co-transformed into Escherichia coli BL21 (DE3) cells. The protein expression was induced by 0.2 mM isopropyl-β-D-thiogalactoside (IPTG) at an OD_600nm_ of 0.6. Cells were growing for 12-16 h at 20℃. The harvested cells were resuspended in the binding buffer (25 mM Tris-HCl pH 7.5, 500 mM NaCl, 3 mM β-mercaptoethanol, 5 mM imidazole, and 1% glycerol). Followed by sonication and centrifugation, the lysate was loaded onto Ni-NTA resin. The resin was washed with binding buffer for three times, and eluted with the binding buffer supplementary with 300 mM imidazole. The eluate was then loaded onto a HiTrap Heparin HP column (Cytiva). Peak fractions containing target protein complex were concentrated and further purified by a Superdex 6 column (Cytiva) equilibrated with the running buffer (25 mM Tris– HCl pH 7.5, 200 mM NaCl, and 2 mM DTT). Peak fractions were collected, and analyzed by SDS-PAGE.

The protein was incubated with target DNA labeled FAM and then loaded onto the fluorescence size exclusion chromatography (FSEC) system^39^ with a Superose 6 column to test the binding efficiency. The absorbance was measured at 492 nm.

### Cryo-EM sample preparation and data collection

To prepare the cryo-EM sample, the protein was concentrated to 1 mg/mL, then Vitrobot Mark IV (FEI) was used to freeze the cryo-EM grids. 3.2 µL samples were applied to the plasma cleaned GIG R1.2/1.3 holey gold grids with the blot force of 2 for 2 s in 100% humidity. The grids were then plunged into the liquid ethane which was cooled by liquid nitrogen. The cryo-EM data was collected using a 300 kV Titan Krios equipped with a Falcon4 direct electron detector (Gatan) with a pixel size of 0.808 or 0.83 Å in a defocus range of -1.2 to -2.5 µm. Each micrograph was dose-fractioned to 32 frames. The total accumulated dose is 60.0 e^-^/Å^2^.

### Protein stability determination

Prior to loading the sample to FSEC^39^, 3 μg proteins were incubated with 0.5 mM metal ions including MgCl_2_, MnCl_2_, CaCl_2_, ZnSO_4_, CoCl_2_, CeCl_2_, and NiCl_2_ on ice for 30 mins, respectively. The samples were loaded to the FSEC system with a Superose 6 column and the absorbance was measured at 280 nm.

### Cryo-EM data analysis

The cryoEM data analysis was performed in cryoSparc^41^ if not noted. For the dataset of Cas5-HNH/Cascade complex unbound to DNA, a total of 7,382 movies were recorded. The beam-induced motion was corrected by MotionCor2^42^. Next, the defocus values were estimated by CTF patch estimation^43^. The protein particles were picked by Blob picker with the minimum and maximum diameters of 100 Å and 220 Å, respectively. After one round of 2D classification, 1,336k good particles were selected from the 1,989k raw particles. To obtain the initial model, six classes were set during the ab-initio reconstruction, the generated models which exhibit good protein features were selected for a further heterogeneous refinement. Notably, the raw particles yielded by Blob picker were used in this round of heterogeneous refinement. Then, four classes were further chosen for another round of heterogeneous refinement. Class1 and class4 were selected for subsequent analysis. For class1, one round of heterogeneous refinement was performed to further remove these junk particles. In order to eliminate the impacts of the preferred orientation, one round of particle rebalance was performed with the rebalance factor of 1.0 following by one round of non-uniform refinement. The retained 20,320 particles were then transformed into Relion^44^ format using Pyem^45^, spIsonet^46^ was finally used to perform the misalignment and anisotropy correction (apo_Conf1). For class4, one round of heterogeneous refinement was performed and the class with the highest resolution was chosen for a final run of non-uniform refinement (apo_Conf2). The resolution for apo_Conf1 and apo_Conf2, as estimated by cryoSparc^41^, were 3.10 Å and 2.47 Å, respectively. DeepEMhancer^47^ was used to sharpen both the maps. For the dataset of Cas5- HNH/Cascade complex bound to target ssDNA, a total of 4,040 micrographs were collected and the data processing is similar to the apo-complex. The final resolution for ssDNA_Conf1 and ssDNA_Conf2 are 3.00 Å and 2.81 Å, respectively.

### Model building

Homology models for Cas5-HNH, Cas6, Cas7, Cas8 and Cas11 were generated by Alphafold^48^. USCF-Chimera^49^ was used to fit the predicted models into the corresponding density map, and all of the models were then combined into one model. The chain IDs were re-assigned in Coot^50^. The model was then manually adjusted in Coot as guided by the high-quality features. The following residues including residues 1-80 in chain C (Cas8), residues 54-138, 148-166 and 314-322 in Chain G (Cas7) and residues 190-208 and 256-263 in chain K (Cas7), were removed due to obscured densities. The structure of RNA was manually built in Coot^50^. Real space refinement in Phenix^51^ was used to refine the model against the cryo-EM map.

We observed significant conformational differences between apo_Conf1 and apo_Conf2, and it is not feasible to simply fit the whole model of apo_Conf1 to the apo_Conf2 cryo-EM map. To overcome this problem, the chains from Conf1 model were saved one by one in Chimera^49^. Next, the saved chains were loaded back to Chimera^49^ and the fitting was performed again into the apo_Conf2 cryo-EM map chain by chain. The chains were combined by Chimera^49^ before proceeding into Coot^50^. Coot was then used to manually adjust the model. Notably, these residues with poor densities including residues 1-80 in chain C (Cas8), residues 54-138 and 195-204 in Chain G (Cas7), residues 91-97 in chain F (Cas7), residues 190-208 and 256-233 in chain K (Cas7) and residues 197-205 in chain J (Cas7) were removed due to the poor densities. Five rounds of real space refinement in Phenix^51^ were performed to refine the model. Data collection and model refinement statistics are concluded in Table S1. A similar procedure was followed to build the model ssDNA_Conf1 and ssDNA_Conf2. Simply, the structure apo_Conf1 was fit to the ssDNA bound maps by Chimera followed by manual refinement in Coot^50^. Phenix^51^ was used to refine the model against the corresponding maps.

### In vitro DNA cleavage assay

The 60-bp target DNA sequence was inserted into a pET-derived vector. FspI (NEB) was used to linearize the resulting plasmids for the target DNA cleavage assay. The Cas5-HNH/Cascade complex and the target DNA substrate were prediluted in a cleavage buffer with 50 mM Tris-HCl, pH 7.5, 100 mM NaCl, 5 mM MgCl2, and 1 mM DTT. To determine the cleavage activity of the Cas5-HNH system, the target DNA was incubated with the WT or the mutants complex at molar ratios ranging from 1:10 to 1:50. The reaction was held at 37℃ for 1 hour. Next, 40 mM EDTA and 1 mg/ml protease K were added to terminate the reaction. A 0.5% TBE agarose gel using StarStain Red Nucleic Acid Dye (GenStar) was used to detect the cleavage product.

### Electrophoretic mobility shift assay (EMSA)

The binding of the target DNA to the Cas5-HNH/Cascade complex was determined using DNA EMSA. The TS and 5’-Cy3 labeled NTS were annealed by heating to 95℃ for 3 minutes followed by progressively cooling to 25℃. The purified Cas5-HNH/Cascade complex was incubated with annealed dsDNA at 37℃ for 30 minutes in a buffer containing 50 mM Tris-HCl, pH 7.5, 10 mM NaCl, and 1 mM DTT, with increasing concentrations of WT or mutant Cas5-HNH complex. All samples were placed onto a 5% native PAGE gel and ran for 30 minutes at 120 V and 4℃. The Tanon 5200 imaging system was used to visualize reaction products.

### Data availability

The coordinates and Cryo-EM maps have been deposited in the Electron Microscopy Data Bank under accession codes EMD-60328 (ssDNA_Conf1), EMD-60330 (ssDNA_Conf2), EMD-60233 (apo_Conf1),EMD-60235 (apo_Conf2), and EMD-60297 (apo_Conf3). The coordinates also have been deposited in the Protein Data Bank under accession codes 8ZP7,8ZP9,8ZLU,8ZM3 and 8ZOL, respectively.

## Acknowledgements

This work was supported by the National Natural Science Foundation of China (E4V4061RA1), Chinese Academy of Sciences (E2VK311RA1) to H.Z., and National Natural Science Foundation of China (32322040 to H.Zhang., and 32300036 to H.Y.). We thank Z.Q.Guo and H.S.Li from Shuimu Biosciences for the cryo-EM data collection. We thank X. Li and Q. He for assistance with experiments.

## Author contributions

Conceptualization: H.Z. and H.Zhu.; Protein purification and Biochemical assays: Q.Z, P.Y. F, L.L.Z.; EM data collection and processing: Y.L., L.W. and H.Zhu.; Structure modelling: Y.L, L.W and H.Zhu.; Supervision: H.Z and H.Zhu.; Manuscript writing: Y.L., L.W., H.Z. and H.Zhu.

