## Supplementary figures for "Structural Basis for RNA-guided DNA degradation by Cas5-HNH/Cascade complex"

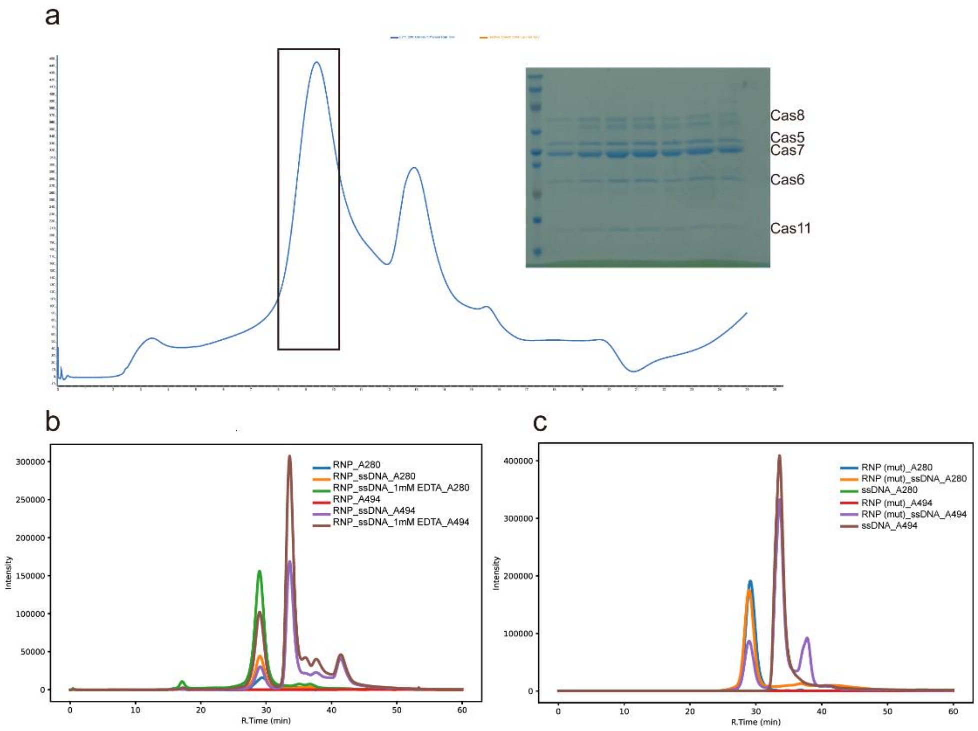


**Supplementary Fig.1 Purification of Cas5-HNH/Cascade complex. a**, The SEC trace showed a mono dispersed sample and the components of the SEC-peak were verified by SDS-PAGE. **b-c,** FSEC showed that both wild-type (**b**) and mutant (**c**) Cas5-HNH/Cascade complex can bound with target ssDNA.

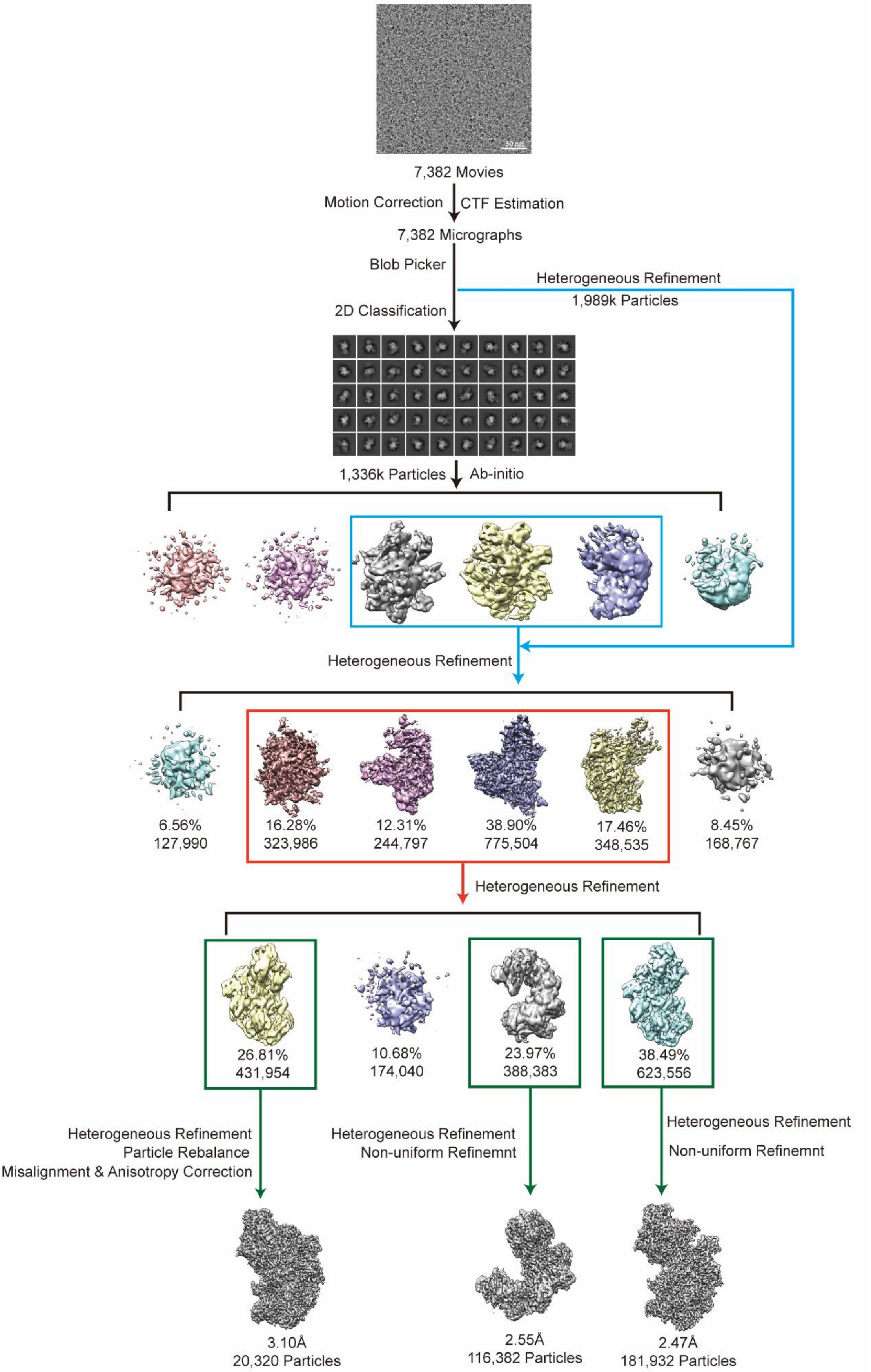


**Supplementary Fig.2 Flow chart for apo-Cas5-HNH/Cascade complex.**

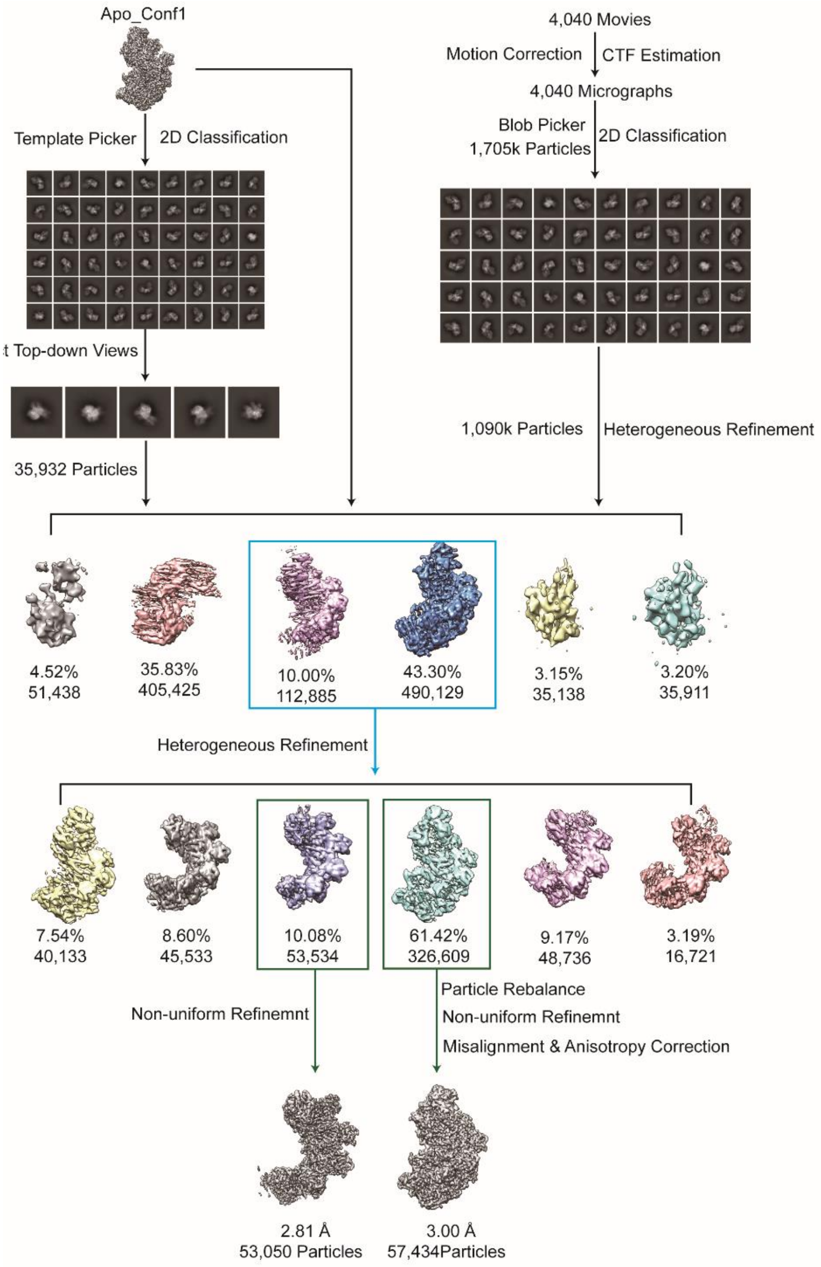


**Supplementary Fig.3 Flow chart for Cas5-HNH/Cascade complex bound with target ssDNA.**

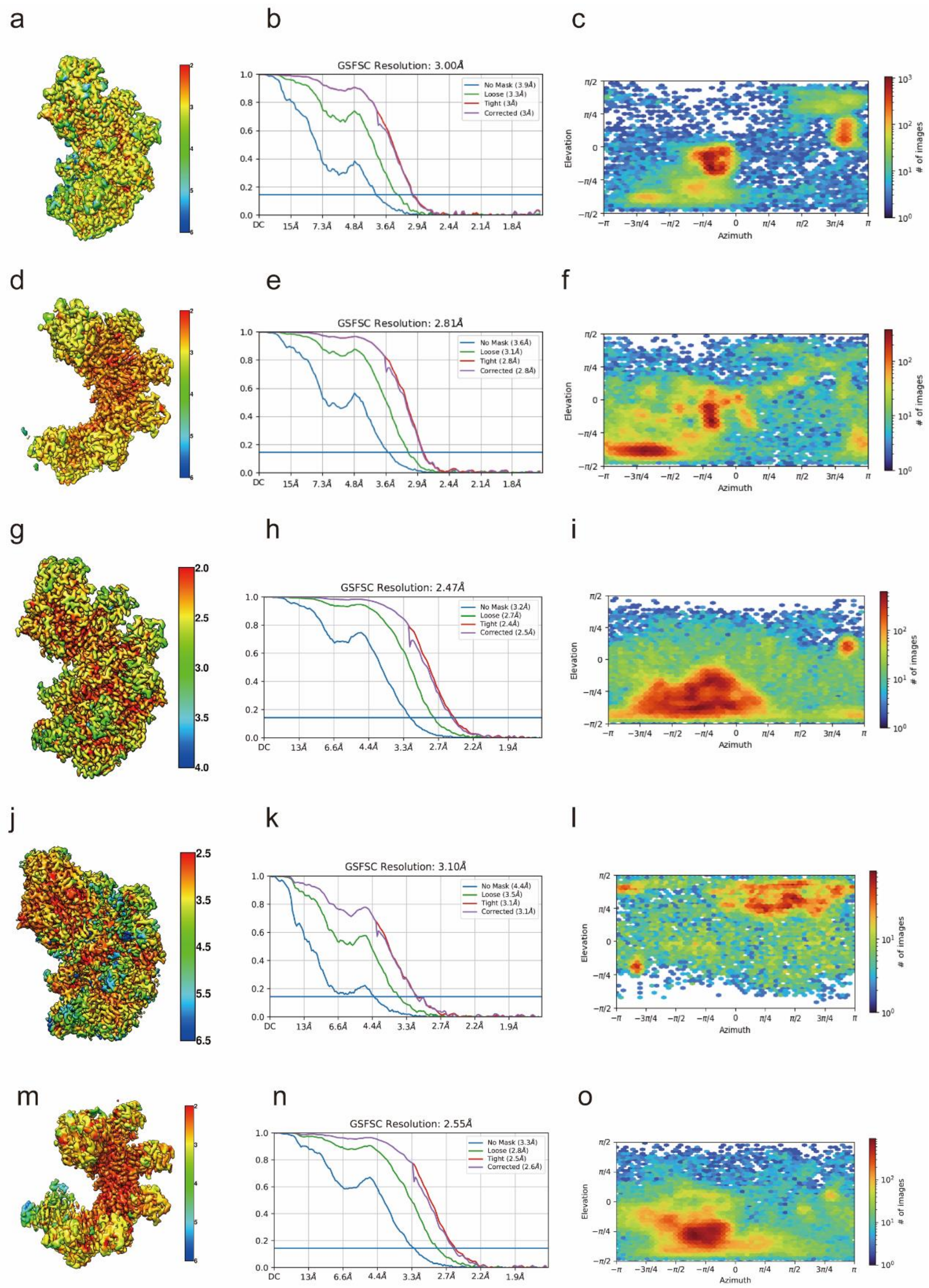


**Supplementary Fig.4 3D reconstruction of the cryo-EM maps.** Local resolution, FSC curves, and particle distributions for intact Cas5-HNH/Cascade complex with target ssDNA (**a-c**), incomplete complex with ssDNA (**d-f**), apo_Conf1(**g-i**), compact apo_Conf2 (**j-l**), and incomplete apo_Conf3 (**m-o**), respectively.

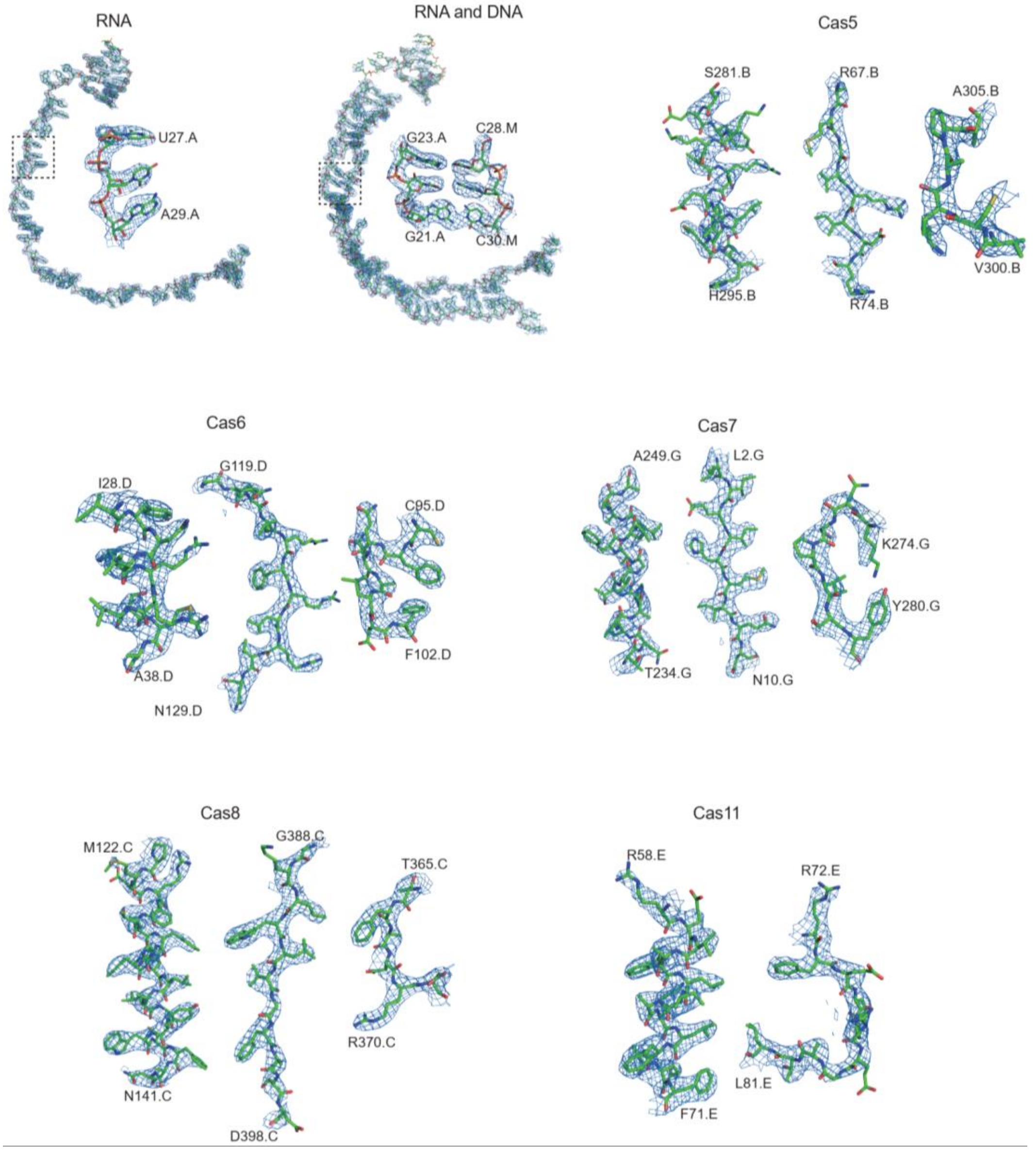


**Supplementary Fig.5 Representative densities of nucleotides and subunits.**

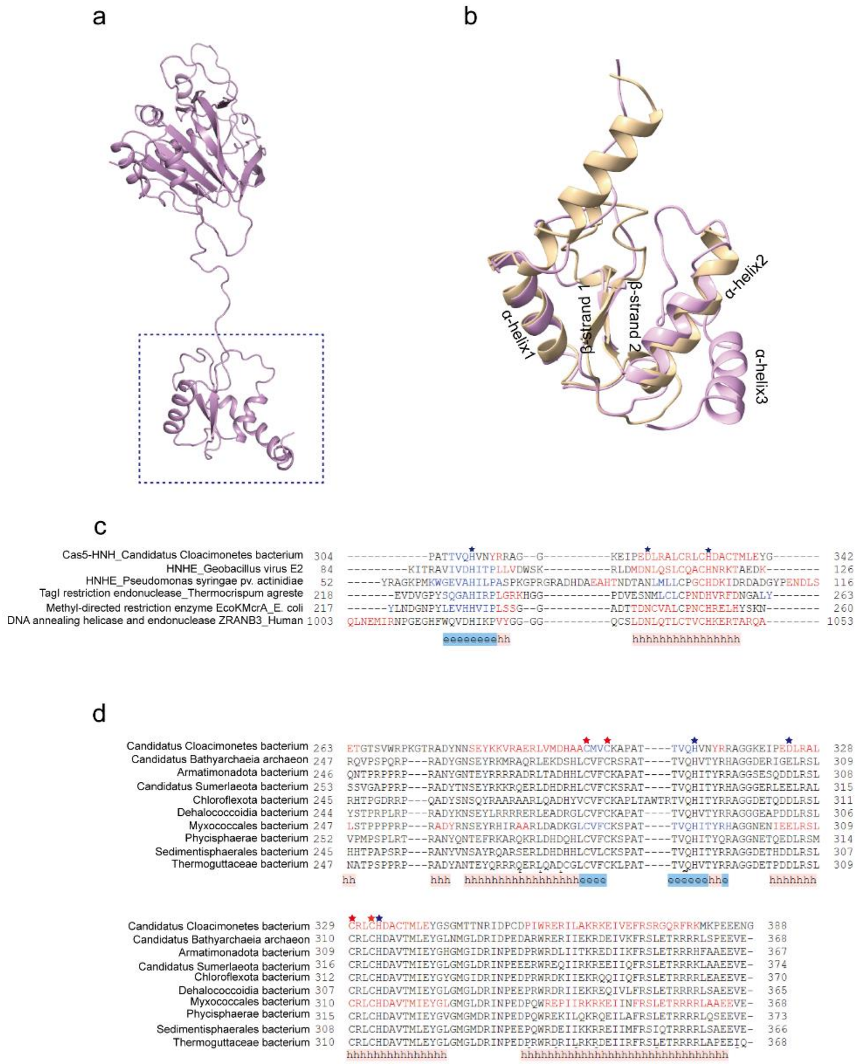


**Supplementary Fig.6 Sequence alignment of HNH domain. a,** Cartoon represent shows the structure of Cas5-HNH subunit. HNH domain is highlighted by a dashed rectangle. **b,** Structural alignment of HNH$^{Cas5}$ (plum) and HNH endonuclease from Geobacillus virus E2 (tan). **c,** Sequence alignment of HNH-containing protein from different species. Featured amino acids are marked by stars. **d,** Sequence alignment of Cascade subunits containing HNH motif. Featured amino acids and four cysteines associated with metal ion-binding are labeled with blue or red stars, respectively.

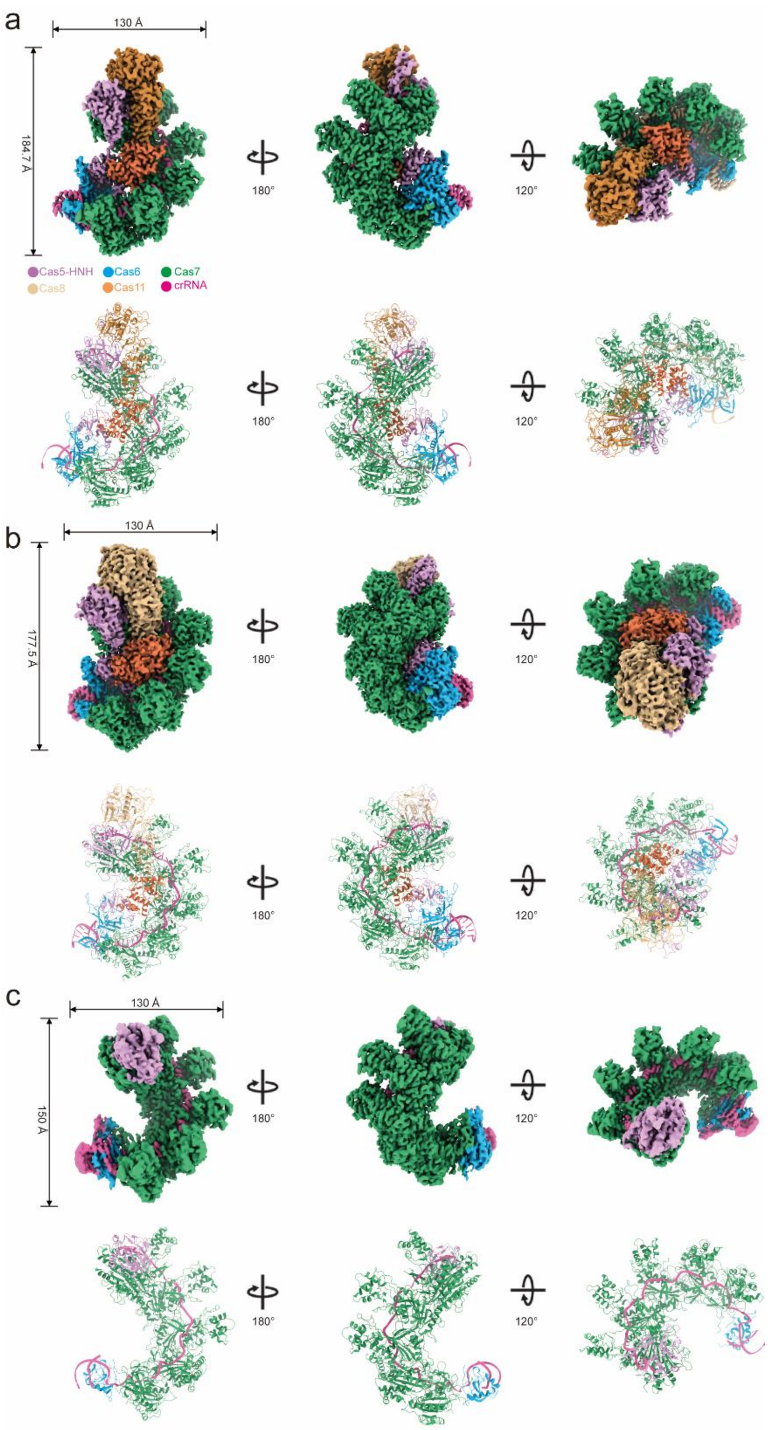


**Supplementary Fig.7 Cryo-EM maps unbound with ssDNA.** Structure of apo_Conf1 (**a**), apo_con2 (**b**) and apo_Conf3 (**c**). The color code is the same to Fig.1.

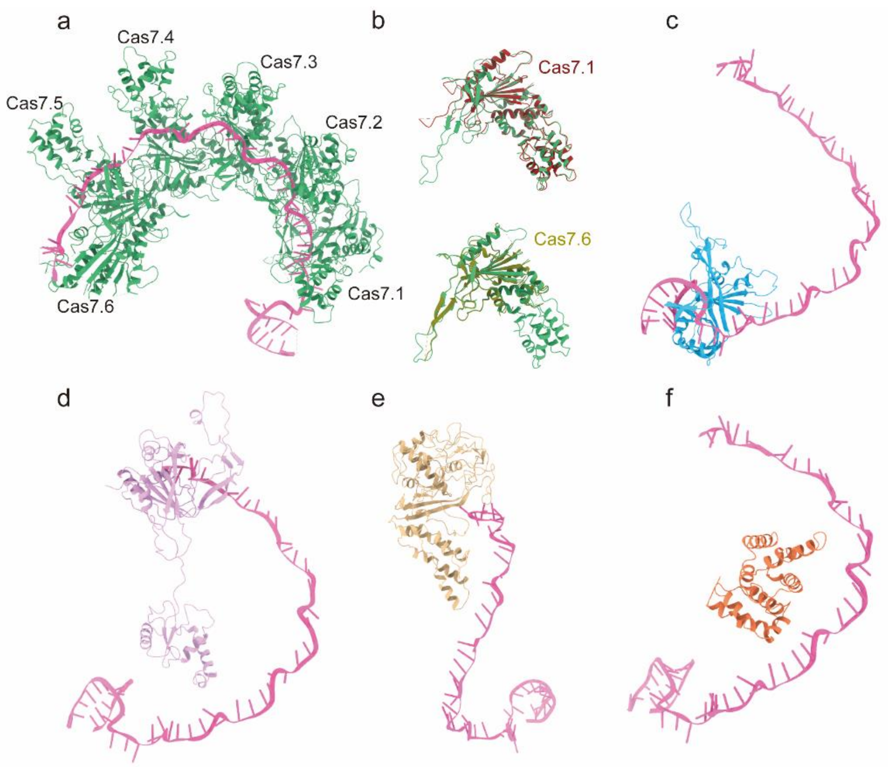


**Supplementary Fig.8 Subunits in Cas5-HNH/Cascade complex. a**, Six Cas7 subunits assemble along with crRNA to form the backbone of complex. **b,** Comparison of classic Cas7 subunit (e.g. Cas7.5) with Cas7.1 (brown) and Cas7.6 (olive),respectively. **c,** Cas6 is clamped by a step-loop architecture of 3'-handle crRNA. **d,** Cas5 caps 5'-handle of crRNA. **e,** Cas8 locates 5'-end of crRNA and it weakly interacts with crRNA. **f,** Cas11 small subunit locates on the center of complex and it has no direct interaction with crRNA.

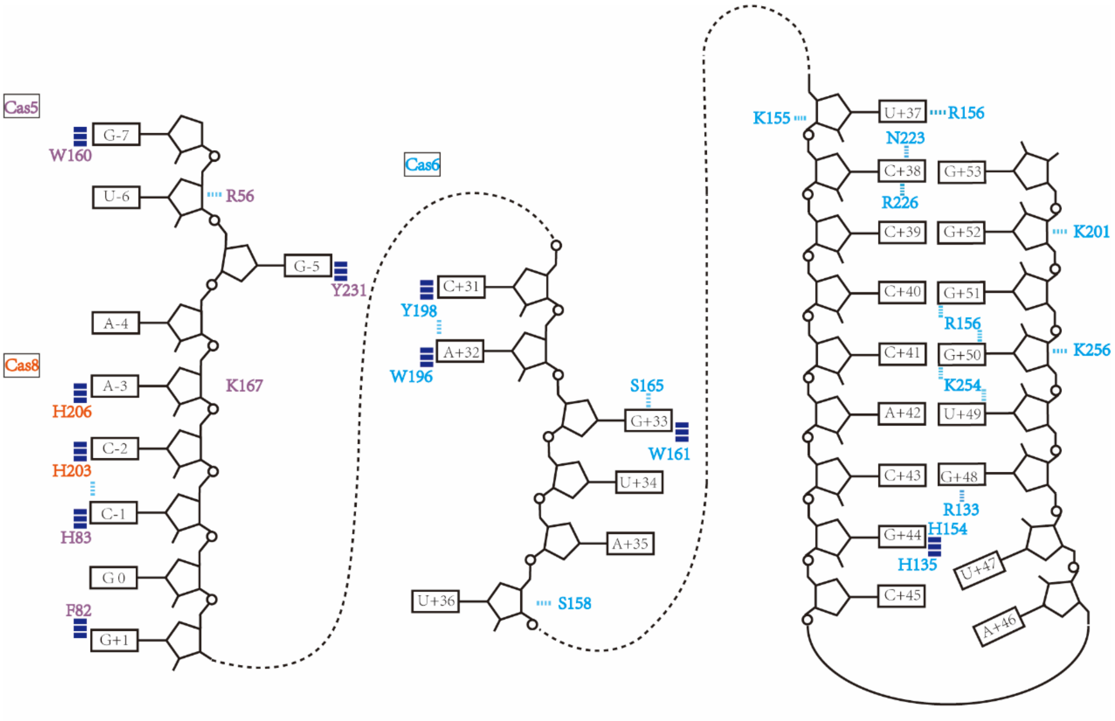


**Supplementary Fig.9 Schematic illustrates of Cas protein-crRNA interactions except for Cas7**. Besides Cas7, RNA associates with Cas5 (plum), Cas6 (cyan), and Cas8 (orange) through hydron bonds (narrow dashed lines) and π-π interaction (rectangles colored by dark blue).

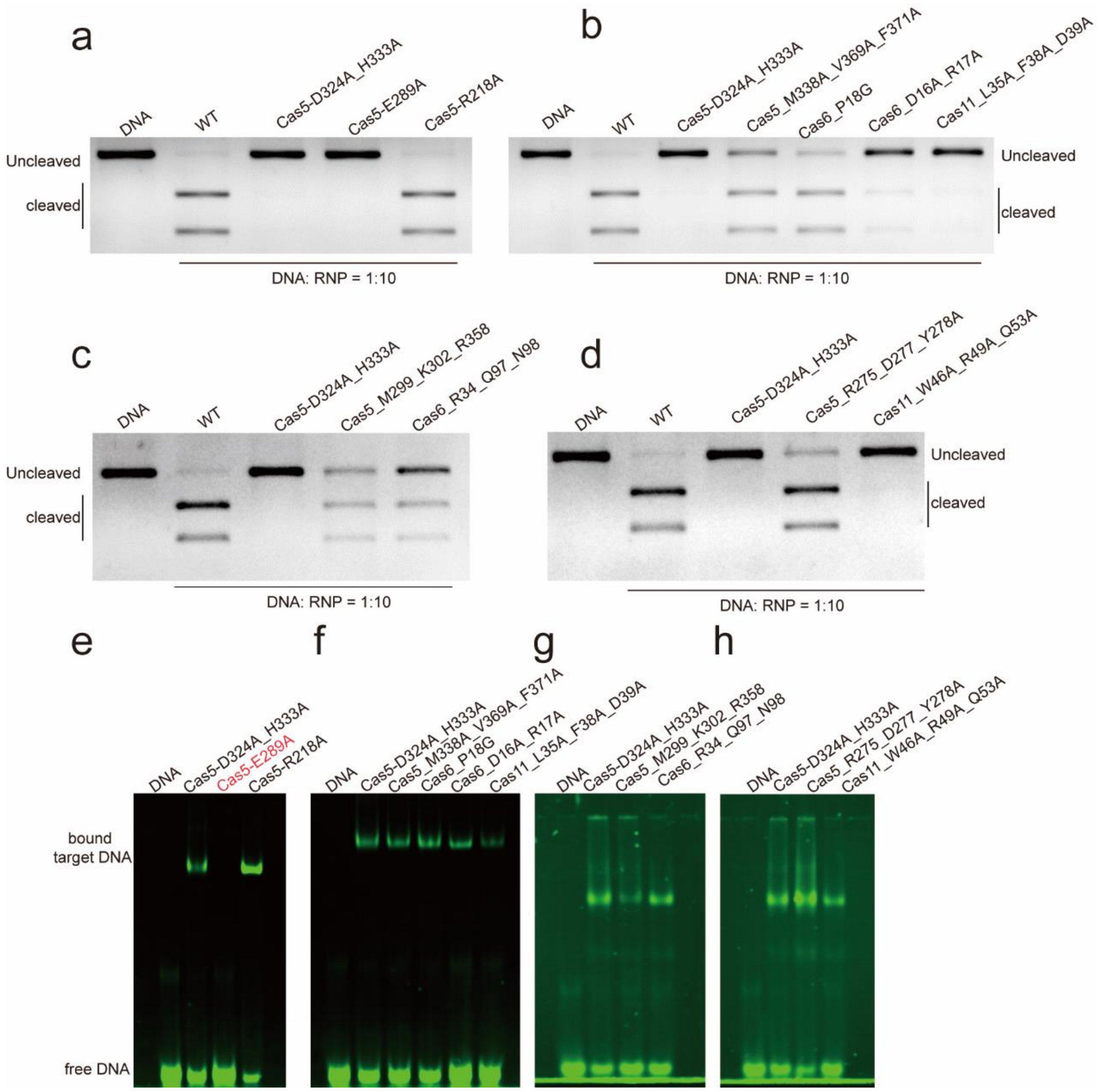


**Supplementary Fig.10 Biochemical analysis for Cas5-HNH/Cascade complex. a-d,** Plasmid digestion analysis indicated that the mutation of residues of Cas5-HNH domain affected the nuclease activity. **e-h,** EMSA results for Cas5-HNH/Cascade complex. The red labeled lane represents the mutant that abolishes the binding of DNA.

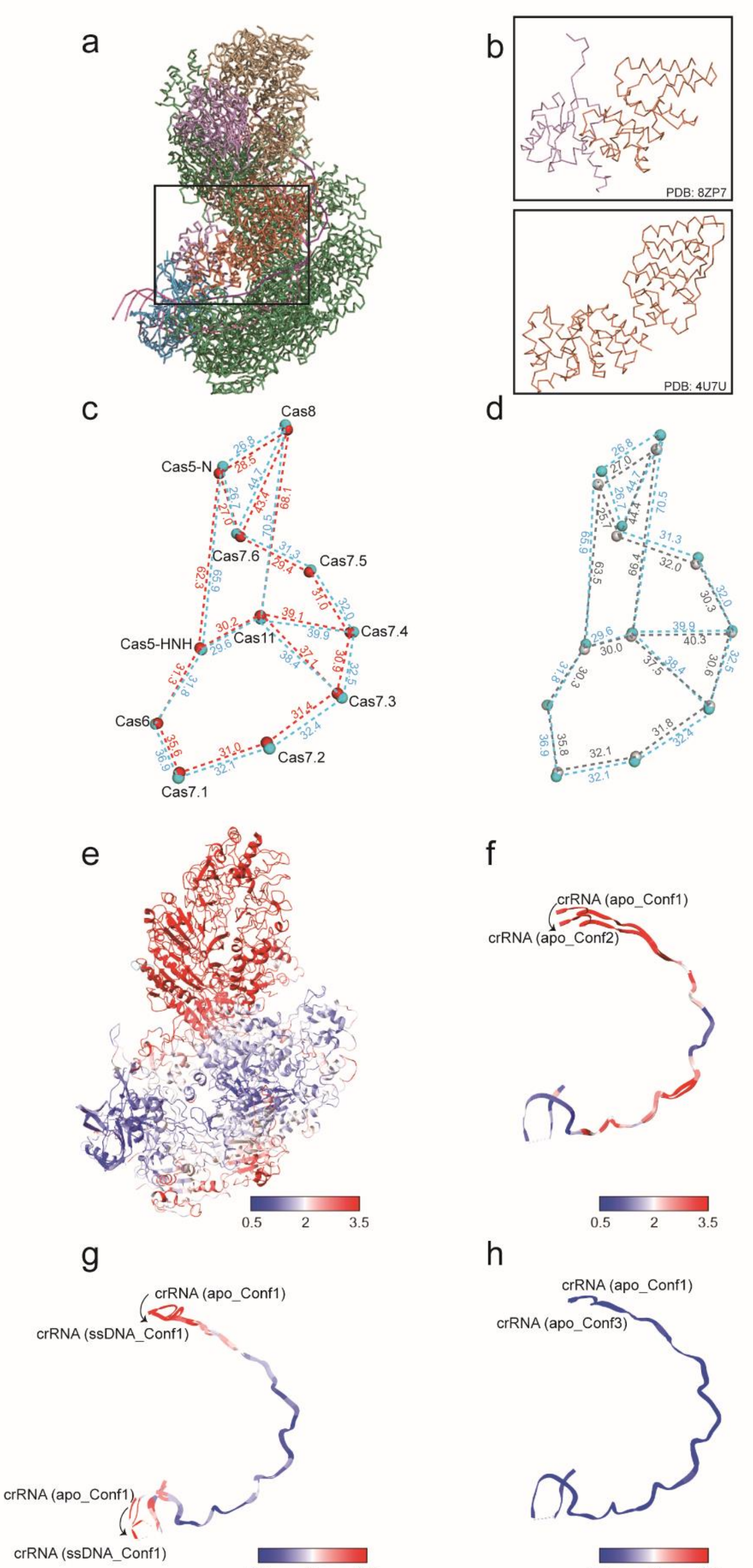


**Supplementary Fig.11 Comparison of Cas5-HNH/Cascade complexes. a,** Comparison of Cas5-HNH/Cascade complex (PDB: 8ZP7) and canonical type I-E Cascade (PDB: 4U7U). **b,** Zoom in views of the main difference of two structures in **(a)**. **c,** Comparison of Cas5-HNH/Cascade complex bound with ssDNA (red) and apo_Conf1(cyan) by center of mass (COM). Distances (Å) between subunits are labeled. **d,** Comparison of apo_Conf1 (cyan) and apo_Conf2 (grey). Distances (Å)

between subunits COM are labeled. **e**, apo_Conf1 was colored by the RMSD value between apo_Conf1 and apo_Conf2 that aligned by crRNA. **f-h,** comparison of the crRNA that was colored by the RMSD value between apo_Conf1 and apo_Conf2 (**f**), between apo_Conf1 and ssDNA_Conf1 (**g**), and between apo_Conf1 and apo_Conf3 (**h**), respectively.

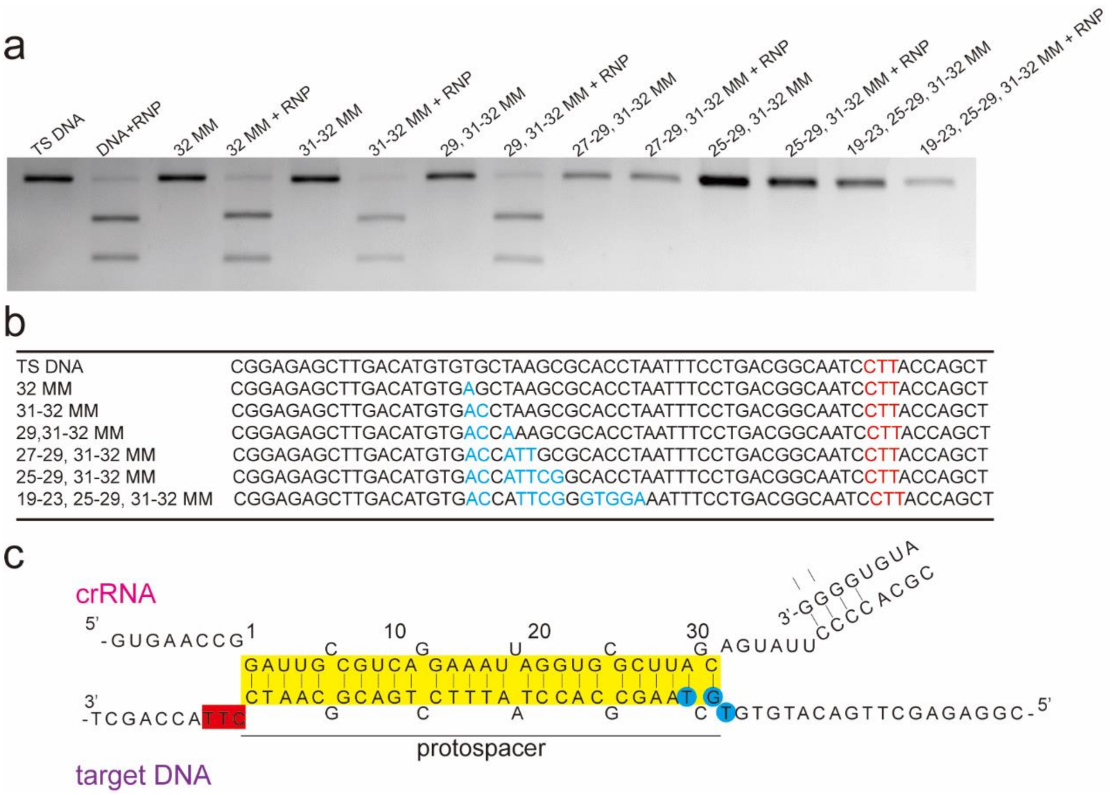


**Supplementary Fig.12 Mismatch tolerance of Cas5-HNH/Cascade complex. a,** In vitro DNA cleavage assay with mismatched substrates (MM: mismatch). **b,** Sequence list of DNA substrates used in a. Mismatch nucleotides are colored in cyan and PAM sequence is noted in red. **c,** Schematic representation of crRNA and the target DNA. The region of spacer is highlighted by yellow rectangle. PAM sequence is showed in red, and mismatches with tolerance by Cas5-HNH/Cascade at PAM-distal positions are labeled with blue circle.

**Supplementary Table 1**

**Cryo-EM data collection, refinement and validation statistics.**

| States | **Apo_Conf1** | **Apo_Conf2** | **Apo_Conf3** | **ssDNA_Conf1** | **ssDNA_Conf2** |
|---|---|---|---|---|---|
| Codes | PDB 8ZLU<br>EMD-60233 | PDB 8ZM3<br>EMD-60235 | PDB 8ZOL<br>EMD-60297 | PDB 8ZP7<br>EMD-60328 | PDB 8ZP9<br>EMD-60330 |
| **Data collection and processing** | | | | | |
| Magnification | | 165,000 | | 96,000 | |
| Voltage (kV) | | 300 | | 300 | |
| Electron exposure (e–/Å$^2$) | | 60 | | 60 | |
| Defocus range (μm) | | -1.8 - -2.5 | | -1.2 - -2.2 | |
| Pixel size (Å) | | 0.83 | | 0.808 | |
| Micrographs (no.) | | 7,382 | | 4,040 | |
| Symmetry imposed | | C1 | | C1 | |
| Total exposure time (s) | | 3.72 | | 7.82 | |
| Detector | | Falcon4 | | Falcon4 | |
| Initial particle images (no.) | | 1,989,579 | | 1,705,634 | |
| Final particle images (no.) | 181,932 | 20,320 | 127,239 | 57,434 | 53,050 |
| Map resolution (Å) | 2.47 | 3.10 | 2.55 | 3.00 | 2.81 |
| FSC threshold | 0.143 | 0.143 | 0.143 | 0.143 | 0.143 |
| Map resolution range (Å) | 2.0-4.0 | 2.5-6.5 | 2.0-6.0 | 2.0-6.0 | 2.0-6.0 |
| *B* factor range (Å$^2$;) | 16.34 - 113.94 | 22.26 - 107.09 | 6.79 - 293.26 | 45.6 - 263.6 | 18.3 - 208.1 |
| **Refinement** | | | | | |
| Initial model used | N/A | N/A | N/A | N/A | N/A |
| Map Correlation Coefficient | 0.81 | 0.75 | 0.70 | 0.85 | 0.79 |
| **Model composition** | | | | | |
| Non-hydrogen atoms | 27,057 | 27,016 | 19,550 | 27,638 | 19,246 |
| Protein residues | 25,799 | 25,758 | 18,292 | 25,711 | 17,760 |
| Ligands | 0 | 0 | 0 | 0 | 0 |
| RNA | 1,258 | 1,258 | 1,258 | 1,258 | 856 |
| DNA | 0 | 0 | 0 | 669 | 630 |
| **R.m.s. deviations** | | | | | |
| Bond lengths (Å) | 0.002 | 0.012 | 0.006 | 0.006 | 0.005 |
| Bond angles (°) | 0.558 | 1.410 | 0.712 | 1.051 | 0.883 |
| **Validation** | | | | | |
| MolProbity score | 2.26 | 2.15 | 1.97 | 2.47 | 2.33 |
| Clashscore | 5.89 | 10.43 | 9.20 | 9.85 | 8.01 |
| Poor rotamers (%) | 4.29 | 0.67 | 0.21 | 3.91 | 3.51 |
| **Ramachandran plot** | | | | | |
| Disallowed (%) | 0.21 | 0.76 | 0.26 | 0.31 | 0.17 |
| Allowed (%) | 6.99 | 11.80 | 7.54 | 8.27 | 7.56 |
| Favored (%) | 92.79 | 87.44 | 92.20 | 91.43 | 92.26 |
